# pH-dependent trapping of cationic amphiphilic drugs perturbs insulin granule homeostasis

**DOI:** 10.64898/2026.02.19.706757

**Authors:** Oleksandra Topcheva, Andreas Müller, Marcello Leomil Zoccoler, Martin Neukam, Pamela Toledo, Katharina Ganß, Anke Sönmez, Carolin Wegbrod, Carla Münster, Klaus-Peter Knoch, Juha Torkko, Sofia Traikov, Michal Grzybek, Michele Solimena

## Abstract

Pancreatic β-cells store insulin in acidic secretory granules (SGs), specialized organelles that also contain monoamine neurotransmitters such as serotonin. Many neuroactive drugs with monoaminergic activity are cationic amphiphilic drugs (CADs) that accumulate in acidic compartments by pH-dependent trapping. Yet, whether insulin SGs represent a site of CAD accumulation and if this affects their properties such as monoamine storage and pH remain unclear.

Here, we show that Slc18a1/VMAT1 is required for vesicular monoamine uptake and maintenance of cellular serotonin levels in insulinoma INS-1 cells. In contrast, neuroactive CADs accumulate via pH-dependent trapping at luminal pH values characteristic of insulin SGs. CADs inhibit VMAT-mediated uptake of the fluorescent monoamine probe FFN206 and induce its efflux to the extracellular space without detectable changes in SG luminal pH. Conversely, natural VMAT substrates such as serotonin and dopamine increase SG pH in a VMAT-dependent manner.

These findings identify insulin SGs as acidic organelles susceptible to CAD accumulation and uncover distinct mechanisms regulating secretory granule homeostasis.

## Introduction

In pancreatic β-cells, insulin is stored in secretory granules (SGs)—acidic organelles with a luminal pH of ∼5.5, maintained by the vacuolar H⁺ -ATPase (vATPase) (Casey *et al*, 2010; Freeman *et al*, 2023; Paroutis *et al*, 2004). This acidic environment is critical for insulin maturation and its crystallization with Zn²⁺ (Aspinwall *et al*, 1997). Beyond insulin, SGs contain several small molecules, including monoamine neurotransmitters (Braun *et al*, 2012; Suckale & Solimena, 2010)

Monoamine neurotransmitters—particularly serotonin (5-HT) and dopamine (DA)—localize to insulin SGs, as demonstrated by early precursor-labeling studies (Ekholm *et al*, 1971; Ericson *et al*, 1977; Lundquist *et al*, 1971) and by glucose-stimulated co-secretion with insulin (Almaca *et al*, 2016; Bennet *et al*, 2015)

Packaging of monoamines into SGs is mediated by vesicular monoamine transporters (Slc18a/VMATs), which function as H⁺ /monoamine antiporters (Liu *et al*, 1992; Pothos *et al*, 1998; Wimalasena, 2011). These transporters display broad substrate permissiveness and interact with a wide range of endogenous monoamines and synthetic compounds (Drew *et al*, 2021; Yelin & Schuldiner, 1995). Two isoforms exist: Slc18a1/ VMAT1 and Slc18a2/VMAT2. In humans, VMAT2 is expressed in pancreatic β-cells (Anlauf *et al*, 2003; Schafer *et al*, 2013), whereas its presence in rodent β-cells remains a matter of controversy (Schafer *et al*., 2013).

Once released from cells, monoamines modulate hormone secretion and islet cell plasticity. Among them, 5-HT has emerged as a key regulator, promoting insulin secretion and β-cell proliferation in states of increased metabolic demand, including pregnancy (Kim *et al*, 2010; Kim *et al*, 2015; Roberts *et al*, 2023). In contrast, DA acts predominantly as an inhibitor of insulin secretion (Aslanoglou *et al*, 2021; Farino *et al*, 2020). Together, these observations support a role of monoamines as modulators of β-cell function and adaptation to metabolic stress (Almaca *et al*., 2016; Ekholm *et al*., 1971; Hamilton *et al*, 2018), with 5-HT representing the best-characterized and physiologically most relevant monoamine in pancreatic β-cells to date.

Alterations in monoaminergic signaling can lead to neuropsychiatric disorders. Both neuropsychiatric disorders and their treatments have been linked to an increased risk of type 2 diabetes (Lee *et al*, 2023; Lindekilde *et al*, 2022). Most widely prescribed antipsychotics and antidepressants are cationic amphiphilic drugs (CADs) (Ballon *et al*, 2014; Barnard *et al*, 2013; Derijks *et al*, 2008). While the effects of CADs on the central nervous system have been extensively studied, potential direct effects of these drugs on pancreatic β-cell function remain incompletely understood (Fathallah *et al*, 2015; Garcia-Tornadu *et al*, 2010; Maines *et al*, 2023).

CADs are lipophilic weak bases characterized by a hydrophobic aromatic core, an ionizable amine group, and high membrane permeability (Funk & Krise, 2012; Kazmi *et al*, 2013). Unlike hydrophilic monoamine neurotransmitters, which require transporters for vesicular uptake, CADs readily diffuse across membranes and become protonated and retained within acidic organelles—a process known as pH-dependent trapping or lysosomotropism (Blumenfeld *et al*, 2024; de Duve *et al*, 1974). While lysosomes represent the best-characterized sites of CAD accumulation, any intracellular compartment with a sufficiently acidic lumen may, in principle, permit such accumulation.

Despite being independent from transporter-facilitated transport, CADs have been reported to inhibit VMAT-mediated uptake due to its pharmacological promiscuity (Hu *et al*, 2013; Stove *et al*, 2022). Another class of monoaminergic drugs, such as the VMAT synthetic substrates amphetamines or MPP⁺, can induce the efflux of neurotransmitters from synaptic vesicles and chromaffin granules and their redistribution into the extracellular space concomitantly with alteration of the vesicular proton gradients (Freyberg *et al*, 2016; Mlinar & Corradetti, 2003; Partilla *et al*, 2006; Rudnick & Wall, 1992a, b; Torres & Ruoho, 2014; Wimalasena *et al*, 2008). Whether similar mechanisms operate in insulin SGs, and how they intersect with pH-dependent trapping of CADs, has not been addressed.

Given the acidic luminal pH of insulin SGs, we hypothesized that they may accumulate neuroactive CADs via pH-dependent trapping. Whether insulin SGs indeed serve as a secondary site of CAD accumulation, and how such accumulation influences monoamine storage and granule properties, has not been systematically examined. Here, we tested whether CAD accumulation in insulin SGs interferes with VMAT function and alters monoamine content or SG pH. Using the rat pancreatic insulinoma INS-1 cell line and VMAT-deficient clones, we combined genetics, imaging, and biochemical approaches to characterize monoamine transport and storage, assess the cellular and subcellular accumulation of the fluorescent VMAT substrate FFN206 and selected neuroactive CADs, and evaluate their effects on monoamine retention and insulin SG pH.

## Results

### VMAT1 is responsible for the monoamine uptake in INS-1 cells

Rat insulinoma INS-1 cells express *Slc18a1* encoding VMAT1. Following the generation of *Slc18a1^−/−^* INS-1 clones (Neukam et al, in preparation), we verified whether VMAT1 is present on insulin SGs by immunostaining these cells with an anti-VMAT1 antibody. As shown in Fig. 1A, VMAT1 immunoreactivity was heterogeneous across INS-1 cells, with VMAT1 detected only in a subpopulation of INS-1 cells and no signal detected in *Slc18a1^−/−^*INS-1 clones (Fig. 1B, C).

**Figure 1.**
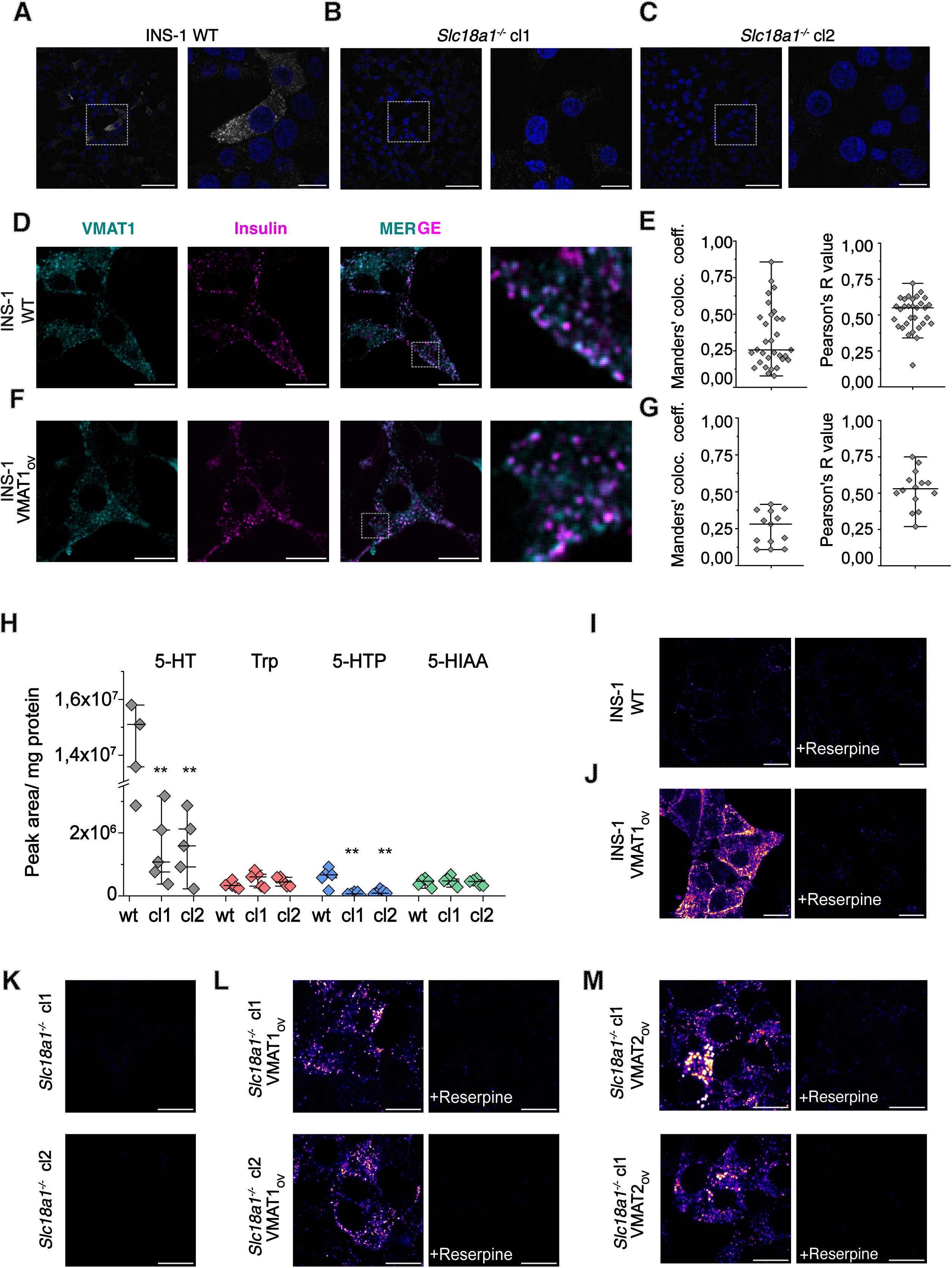
Characterization of monoamine transport in INS-1 cells. Immunostaining of (A) INS-1 and (B, C) *Slc18a1^−/−^* clones with anti-VMAT1 antibody. The staining of INS-1 cells suggests differences in the expression levels of VMAT1 among cells. (D, F) Immunostaining and (E, G) colocalization analysis of insulin and (D) endogenous VMAT1 expression or (F) upon its overexpression (VMAT1*_OV_*). The overexpressed protein shows a similar distribution pattern to the endogenous protein (different acquisition gain and laser power). Co-localization analysis of VMAT1 and insulin in WT (n=3, total 31 fields of view), and VMAT1*_OV_* (n=2, total 13 fields of view). (H) LC MS/MS analysis of endogenous content of serotonin (5-HT), tryptophane (Trp), 5-hydroxytryptophane (5-HTP) and 5-hydroxyindoleacetic acid (5-HIAA) in INS-1. (I) Accumulation of FFN206 in INS-1 cells with endogenous expression of VMAT1 and (J) upon its overexpression. The cells were supplied with 2 μM FFN206 for 60 min. FFN206 accumulation is inhibited with 2 μM reserpine. (K) *Slc18a1^−/−^* INS1 clones incubated in the same conditions with FFN206, showed no accumulation of the compound. Overexpression of either (L) VMAT1 or (M) VMAT2 in *Slc18a1^−/−^*INS1 rescued the uptake of FFN206, which was blocked by 2 μM reserpine. Scale bar for overview images on (A) is 50 μm; for the remaining images 10 μm.

For subsequent analyses, cells displaying higher VMAT1 immunoreactivity were selected for the analysis of endogenous VMAT1 localization. Confocal imaging of INS-1 cells immunostained with antibodies against VMAT1 and insulin showed partial co-localization (Fig. 1D) with a median Manders’ colocalization coefficient of 25.2% (Insulin to VMAT1), indicating partial but not exclusive overlap with insulin-positive SGs (Fig. 1E). A similar colocalization pattern was observed upon the overexpression of rat VMAT1 (Fig.1F). In this case, the Manders’ colocalization coefficient of the immunoreactivity signals for VMAT1 and insulin was 28.2% (Fig. 1G). Similar results were observed using structured illumination microscopy (SIM), where VMAT1 showed a more dispersed intracellular distribution, with signal detected beyond secretory granules (SGs). In addition, not all SGs were positive for VMAT1 (EV1A).

Despite the detection of a weak diffused fluorescent signal in both *Slc18a1^−/−^* clones, the immunoreactivity against the VMAT1 antibody was considerably lower than in wild-type (WT) INS-1 cells (Fig. 1A-C). Incubation with the peptide used to raise the anti-VMAT1 antibody blocked its immunoreactivity, thus corroborating the specificity of the immunostaining (EV1B).

Since VMAT1 can transport all monoamine neurotransmitters (Liu *et al*., 1992; Pothos *et al*., 1998; Wimalasena, 2011) we assessed if any of these are endogenously present in INS-1 cells and if genetic deletion of *Slc18a1* has any effect on their intracellular levels.

Using LC-MS/MS, we detected 5-HT and its precursors, 5-hydroxytryptophan (5-HTP) and tryptophan (Trp), as well as its metabolic product, 5-hydroxyindoleacetic acid (5-HIAA) in both WT and *Slc18a1^−/−^*INS1 clones (Fig. 1H). *Slc18a1* deletion was associated with a significant reduction in 5-HT and 5-HTP levels, although to different extents, but did not affect Trp or 5-HIAA levels (Fig. 1H). We further verified that the reduction of 5-HT did not result from decreased expression of its biosynthetic enzymes Aadc and Tph1. Immunoblot analysis showed that the levels of these enzymes in *Slc18a1 ^−/−^* clones were unchanged or even increased compared to WT INS-1 cells (EV1C). Furthermore, VMAT1 overexpression (VMAT1*_OV_*) in both clones rescued 5-HT and 5-HTP levels, consistent with a role of VMAT1 in monoamine accumulation in INS-1 cells (EV 1D).

We next employed a functional readout for the monoamine uptake in living cells using the False Fluorescence Neurotransmitter (FFN206) - a probe that mimics the natural substrates of VMATs (Hu *et al*., 2013). FFN206 has been used to study the activity of VMAT2 in mammalian cells or in synaptic terminals of the fly brain (Freyberg *et al*., 2016; Hu *et al*., 2013), but to our knowledge has not been applied to pancreatic β cells. After establishing the optimal working concentration for FFN206 imaging in INS-1 cells (1–5 µM; EV1E, F), INS-1 cells, VMAT1_OV_ and *Slc18a1⁻ /⁻* INS-1 cells were incubated with 2 µM FFN206 with or without 2 µM reserpine, a high-affinity VMAT inhibitor (Fig. 1I-M).

*Slc18a1* deletion strongly reduced FFN206 uptake in both clones (Fig. 1K), while VMAT1 overexpression restored FFN206 uptake in a reserpine-sensitive fashion (Fig.1L). As observed previously, VMAT1 overexpression enhanced FFN206 accumulation in intracellular structures compared to WT INS-1 cells (Fig. 1I, J). The overall pattern of FFN206 accumulation was spatially similar but increased in intensity between WT and VMAT1_OV_ cells, supporting comparable subcellular localization (Fig. 1I, J). VMAT2 overexpression also rescued FFN206 uptake in Slc18a1^⁻ /⁻^ INS-1 cells; however, FFN206-positive structures appeared larger and more heterogeneous (Fig. 1M). Therefore, subsequent analyses were conducted in VMAT1_OV_ cells.

Since the uptake of FFN206 is VMAT-dependent, its fluorescence likely corresponds to the subcellular localization of VMAT1, and therefore is detected in the same intracellular compartments where monoamine transport occurs, which to some extents are the SGs (Fig.1D-G).

### Cells accumulate both synthetic neuroactive weak bases and natural neurotransmitters by VMAT-(in)dependent mechanisms

Monoamine neurotransmitters are organic weak bases with limited membrane permeability (EV Table1). Thus, they rely on transporter-mediated uptake and vesicular sequestration (Lawal & Krantz, 2013; Norregaard & Gether, 2001; Torres *et al*, 2003; Vieira & Wang, 2021). In contrast, cationic amphiphilic drugs (CADs) readily permeate membranes and accumulate in acidic organelles (Blumenfeld *et al*., 2024; Funk & Krise, 2012; Kazmi *et al*., 2013).

To investigate if CADs accumulate in insulin SGs of pancreatic β cells, driven by their pH, representative molecules from different drug classes were selected and incubated with INS-1 cells. These selected compounds share some key features. First, they are all membrane-permeable weak bases, as reflected by their LogP and pKa (EV Table2), and second, they represent drug classes associated with increased diabetes risk (Lee *et al*., 2023; Lindekilde *et al*., 2022).

For the initial assessment a classical model CAD, chloroquine (CQ) was selected. It has an intrinsic fluorescence; thus, it is convenient to visualize on thin-layer chromatography plates. Comparatively, other CAD molecules have lower detection sensitivity (EV2). CQ accumulation was compared to VMAT substrates: FFN206 and 5-HT (supplied as 5-HTP), both of which also exhibit detectable fluorescence under UV irradiation (EV2A).

TLC analysis from INS-1 extracts showed that transporter expression did not influence CQ accumulation, as intracellular levels were similar in VMAT1-expressing and non-expressing cells (Fig. 2A, B). In contrast, both 5-HT and FFN206 accumulated only in VMAT1-expressing cells (Fig. 2A, B).

**Figure 2.**
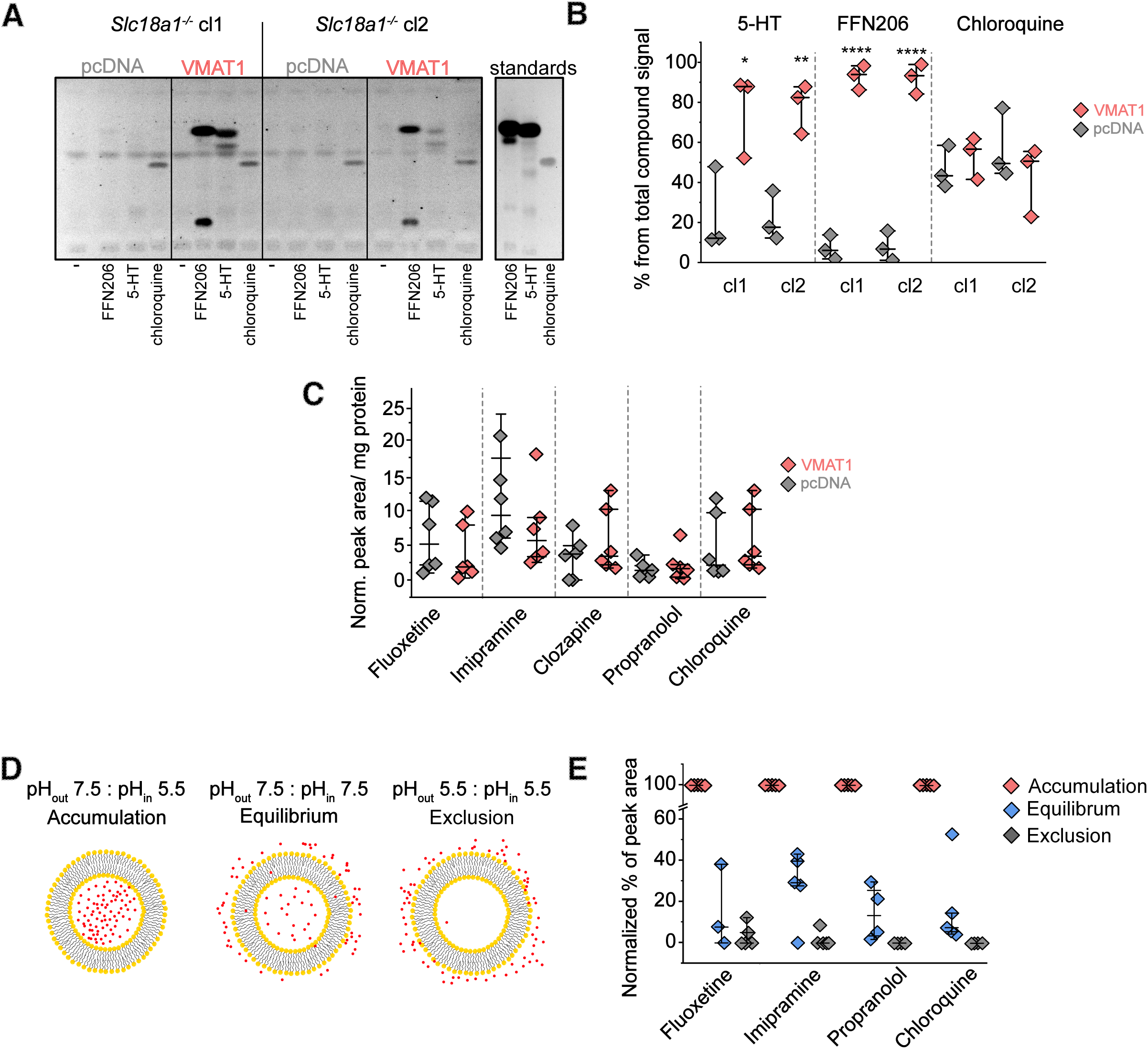
Accumulation of CADs in acidic compartments of beta cells. (A) TLC analysis of Chloroquine, 5-HT, and FFN206 accumulation in *Slc18a1^−/−^* INS1 clone 1 and clone 2, transfected with either an empty vector control (pcDNA) or a rescue construct (VMAT1). Cells were supplied with one of the following 20 µM of 5-HTP (precursor of 5-HT), 2 µM of FFN206 or 20 µM chloroquine. (B) Quantification of TLC (n=3). (C) *Slc18a1^−/−^* NS1 clone 1 and clone 2 transfected with empty vector control (pcDNA) or a rescue construct (VMAT1) were supplied for 60 min with one of the following: 1 µM Fluoxetine, 1 µM Imipramine, 1 µM Clozapine, 1 µM Propranolol, 1 µM Chloroquine. The uptake of these compounds was analyzed using LC-MS/MS. The plot shows pooled data from n=6 (n=3 for each *Slc18a1^−/−^*INS1 clone independently). (D) Schematic representation of pH-dependent accumulation of CADs in liposomes and (E) Quantification of CADs accumulation in these, using MS (n=5; Propranolol n=4). Data normalized by min/max*100%

Hence, while cellular accumulation of 5-HT and FFN206 probably involves compartmentalization in VMAT-containing organelles, CQ accumulates in cells based on its physicochemical properties (Blumenfeld *et al*., 2024; de Duve *et al*., 1974).

The cellular accumulation of other CADs was further measured with LC MS/MS. With an improved detection sensitivity LC MS/MS allowed the analysis of compounds that otherwise would not be detected on TLC. It additionally allowed the use of lower drug concentrations (1 µM final). Results showed that other CADs: Fluoxetine, Propranolol, Imipramine and Clozapine, as well as lower concentration of CQ efficiently accumulated in the cells. This effect was VMAT-independent (Fig. 2C).

Acidic pH of lysosomes drives CADs accumulation, which is an acknowledged phenomenon (Funk & Krise, 2012; Kazmi *et al*., 2013). In contrast to lysosomes, SG have less acidic pH of ∼5.5 (Colomer *et al*, 1996; Davidson *et al*, 1987; De Lorenzi *et al*, 2025). To estimate if SG’s pH still enables CAD accumulation, we prepared liposomes with defined luminal (internal) and external pH (Fig. 2D).

The internal pH was adjusted to either an acidic pH of 5.5, mimicking the pH of insulin SGs, or a neutral pH of 7.5, to account for drug membrane permeability. Before the experiment, all external buffers were exchanged to create models with or without pH gradients (Fig. 2D). Resulting liposomes were then incubated with CADs, and after pelleting, drug accumulation was estimated using MS (EV3).

As a result, liposomes with a pH gradient (pH 5.5 inside, pH 7.5 outside) exhibited highest drug content (accumulation) (Fig. 2E). In comparison, liposomes with a neutral pH (7.5 for internal and external pH) showed lower signals and liposomes with both acidic (5.5) internal and external pH showed negligible signal. The latter possibly accounts for CAD protonation already in the external buffer, preventing membrane permeation (Fig. 2E).

Together, these results indicate that intracellular compartments with a luminal pH of ∼5.5, such as insulin SGs, can act as sites of CAD accumulation.

### CADs inhibit FFN206 accumulation and induce its efflux outside the cell

Because most of the selected CADs are monoaminergic drugs, we tested whether they affect VMAT-mediated FFN206 uptake.

The cells treated with CADs at different concentrations were stained with FFN206 to monitor its uptake. For the assessment, confocal microscopy in VMAT1_OV_ INS-1 cells and a plate reader assay with *Slc18a1*^−/−^ clones were used. Both assays indicated that CAD treatment inhibited FFN206 uptake in a concentration-dependent manner (Fig. 3A, B).

**Figure 3.**
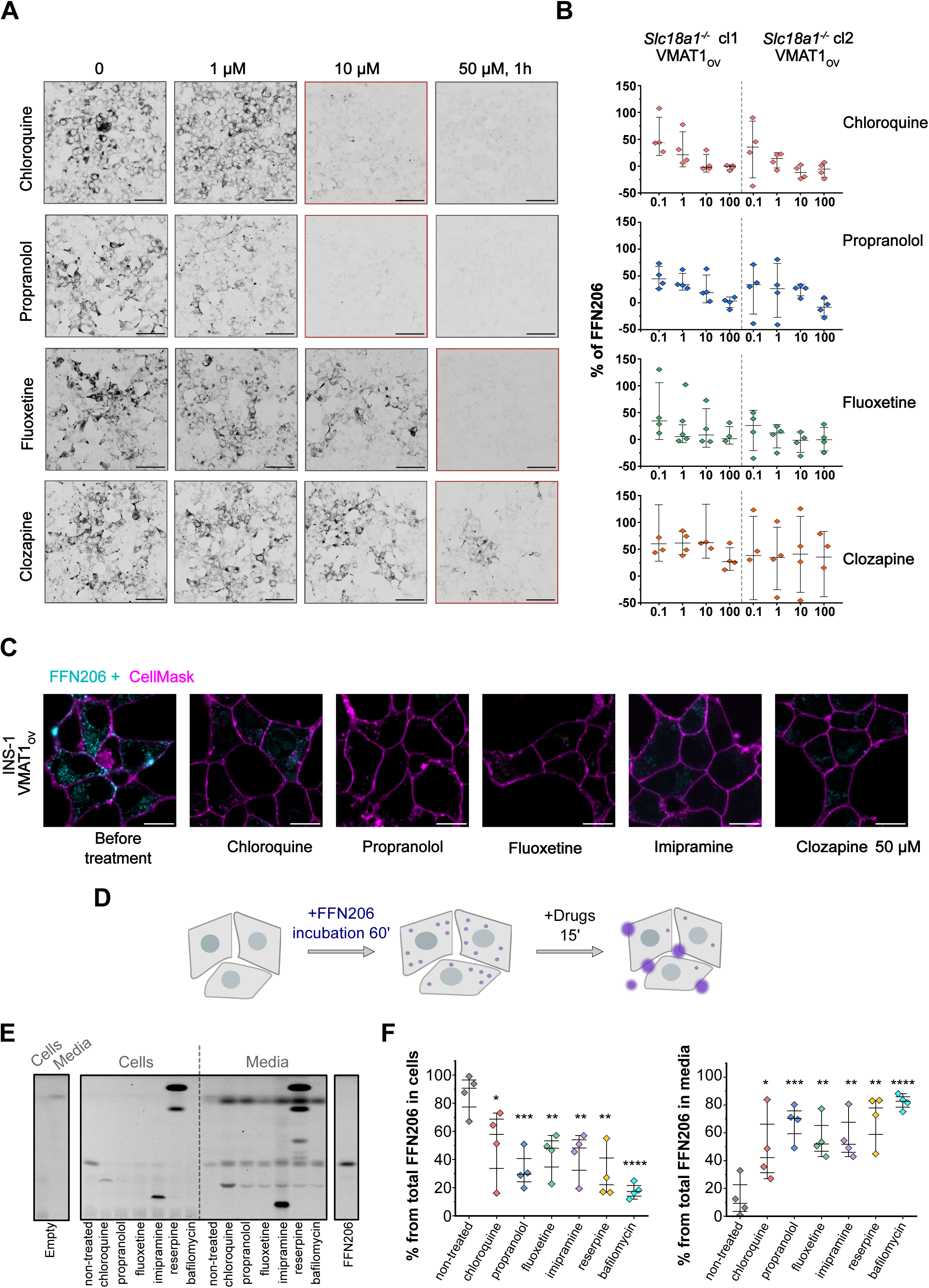
Effects of CAD treatment on FFN206 accumulation and retention. CADs inhibit FFN206 uptake as assayed by FFN206 fluorescence using (A) confocal microscopy (representative images; n=3) or (B) a 96-well plate format (n=4). *Slc18a1^−/−^*INS-1 cells VMAT1*_OV_* were incubated with 2μM FFN206 following treatment with increasing concentrations of various CADs. As a background *Slc18a1^−/−^* INS-1 cells transfected with empty vector (pcDNA) and supplied with 2 μM FFN206 were used. The non treated sample was taken as 100% of FFN206 signal. (C) Confocal images showing INS-1 cells VMAT1_OV_ preincubated with 2μM FFN206 and then acutely treated with various CADs, which leads to a loss of the FFN206 fluorescence. (D) Schematic representation of the experiment (E-F). (E) TLC analysis of the FFN206 extracted from the cells or 500 µL of the growth media after treatment with various CADs and (F) quantification (n=4). Scale bar for confocal microscopy (A) 50 µm) and (C) 10 µm.

To determine whether CADs could, besides inhibition, induce efflux of pre-accumulated FFN206, VMAT1_OV_ INS-1 cells were loaded with FFN206 and subsequently treated with various CADs. After treatment, cells were immediately imaged by confocal microscopy. This resulted in a pronounced loss of FFN206 staining (Fig. 3C).

Next, we approached the FFN206 efflux from INS-1 cells by TLC analysis, which allowed analysis of bulk cellular content. First, we identified the minimal effective efflux conditions with VMAT1_OV_ INS-1 cells loaded with FFN206 which were subsequently incubated with increasing CAD concentrations (EV4A-D). Alternative mechanisms, such as pH dissipation with vATPase inhibitor bafilomycin A (BafA) or inhibition of reuptake with a potent VMAT inhibitor-reserpine, were addressed (EV4E). Concluding the setup, further experiments were conducted with imipramine replacing clozapine since the latter showed greater efflux efficiency (EV4C, D).

To further investigate whether the cellular loss of FFN206 occurs due to its efflux outside the cell, both cellular content and media were analyzed, following the treatment. TLC-based analysis showed that CAD treatment reduced cellular FFN206 signal while increasing FFN206 in the extracellular medium (Fig. 3D-F). This efflux varied and showed between 32,8% and 61% median reduction depending on the CAD. The greatest efflux was reached upon the pH dissipation with BafA with a median 72,4%.

Collectively, these data show that CADs, VMAT inhibitor reserpine, as well as pH dissipation by BafA, lead to the redistribution of previously accumulated FFN206 to the extracellular space.

### CADs, in contrast to VMAT natural substrates, do not impact SG pH

To determine whether FFN206 efflux was associated with SG pH changes, SG pH was measured using FLIM of a granule-targeted pH reporter ICA512-Resp18HD-eCFP (Neukam *et al*, 2017)(preprint).

First, it was determined if VMAT1 overexpression *per se* affected the pH of SGs. *Slc18a1*^−/−^ INS1 clones were transfected with either VMAT1 or an empty vector (pcDNA).

In one of the two clones (cl1), VMAT1 overexpression led to a noticeable difference in the luminal pH (Fig. 4A, B). However, such an effect was not observed for the other clone (cl2). Therefore, VMAT1 expression *per se* did not consistently alter SG pH across *Slc18a1^−/−^*clones (within the resolution of the reporter and calibration plateau (*flim_granules_analysis*; GitHub).

**Figure 4.**
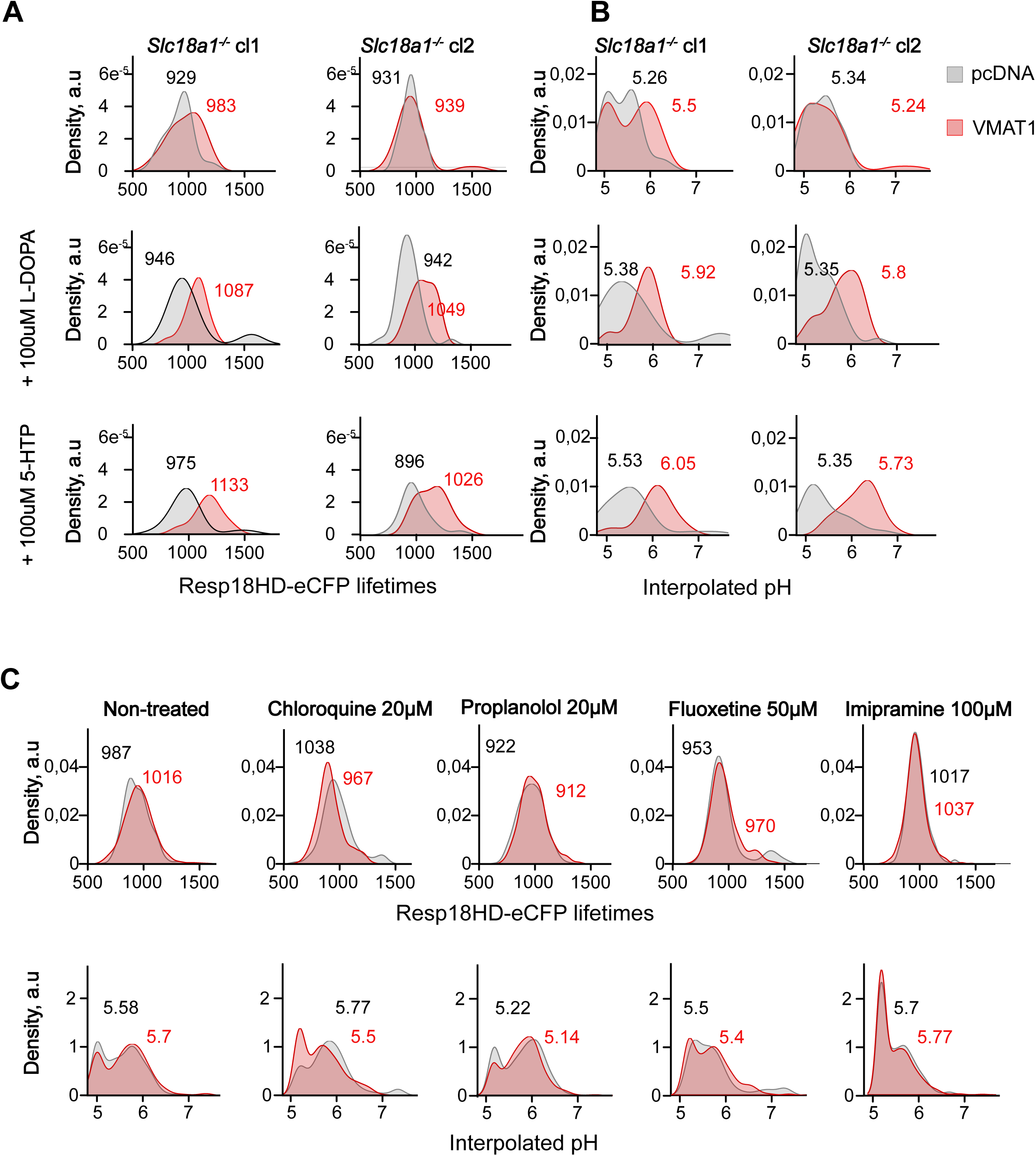
Influence of 5HTP, LDOPA or CADs on the pH of SG. Insulin SG pH measurements of *Slc18a1^−/−^* clones expressing the granular pH reporter ICA512-RESP18HD-eCFP and transfected with either VMAT1 or empty vector (pcDNA). (A) Plots depict the eCFP lifetime distribution of 300 random instances from n=5 biological replicates for each *Slc18a1^−/−^* INS1 clone (cl1 and cl2) or (B) interpolated pH values. Cells treated with either 100 µM 5-HTP or 100 µM LDOPA show shift of SG pH values. (C) CADs’ treatments did not change lifetime distribution or pH. 150 random instances from n = 4 biological replicates pooled from both clones were used for quantification. The numbers on all the plots corresponds to the median lifetime or pH of all the measured instances.

Considering that VMAT substrates were shown to increase synaptic vesicles pH (Freyberg *et al*., 2016), we aimed to verify if natural neurotransmitters serotonin and dopamine can raise the SG pH. Treatment with serotonin and dopamine precursors (5-HTP or L-DOPA) raised the SG pH in a VMAT-dependent way (Fig. 4A, B).

Finally, the influence of CADs on the SG pH was tested. Under conditions that induced FFN206 efflux (Fig. 3D-F), CADs did not change the SG pH. Even CQ, which is known to raise pH of intracellular organelles (Maxfield, 1982; Ohkuma & Poole, 1978; Poole & Ohkuma, 1981), did not measurably alter SG pH under these conditions (Fig. 4C).

Thus, although all of the CADs induced the FFN206 efflux (Fig. 3C-F), they did not alter the pH of SGs (Fig. 4C). Meanwhile, direct precursors of monoamine neurotransmitters raised the SG pH via a VMAT-dependent mechanism (Fig. 4A, B).

### FFN206 does not mimic 5-HT in terms of induced efflux

FFN206 provides a valuable tool for visualizing VMAT-mediated uptake (Hu *et al*., 2013; Yaffe *et al*, 2016). However, it is unclear to what extent it mimics the naturally occurring monoamines. To compare the behavior of FFN206 to that of 5-HT, VMAT1_OV_ INS-1 cells were loaded with 5-HT (via 5-HTP) (Fig. 5A, B). To verify that the setup was functioning as expected, cells were preloaded with FFN206 and subsequently treated with fluoxetine, reserpine and BafA (Fig. 5C-D).

**Fig 5.**
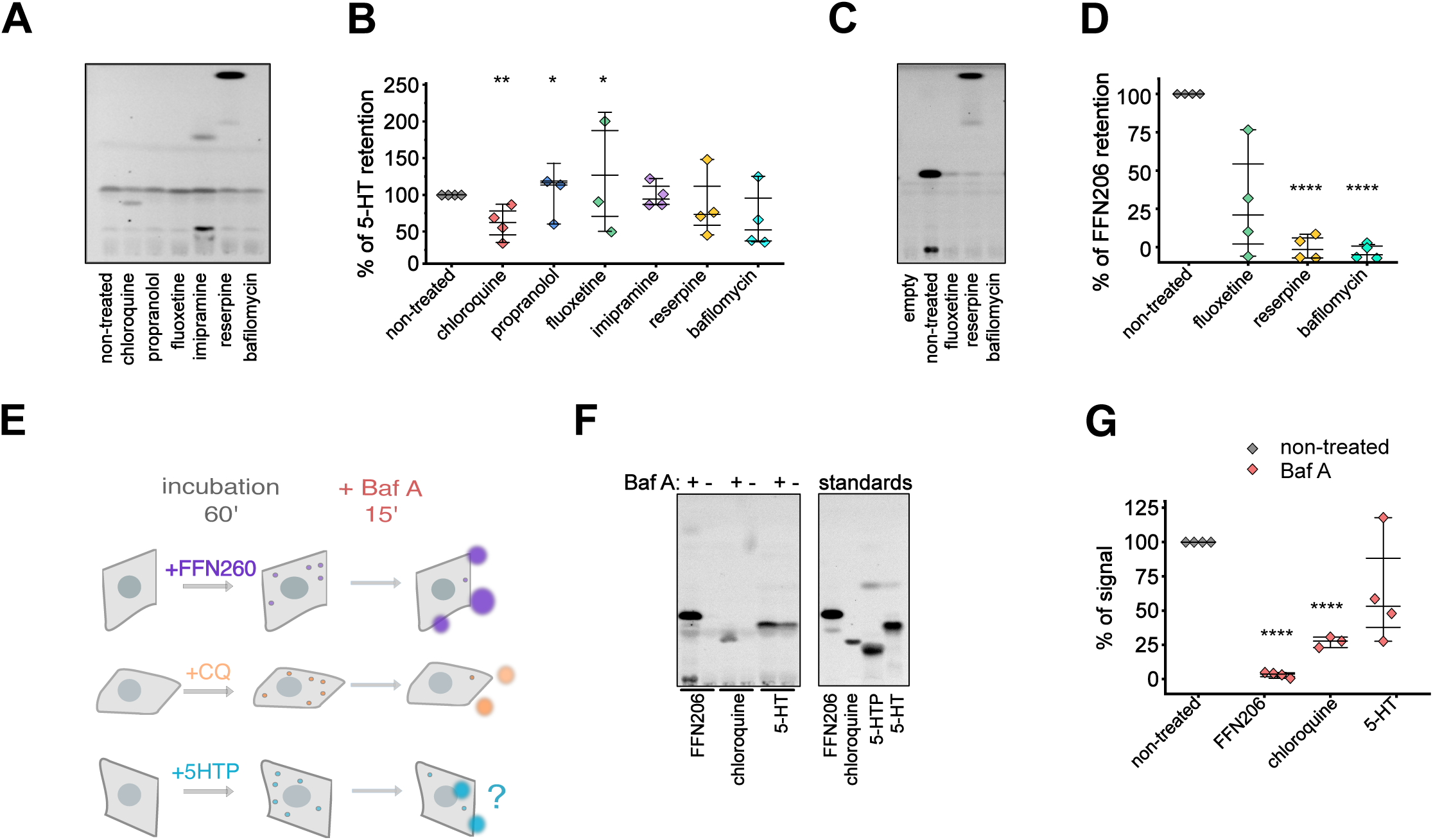
Comparison of the efflux of 5-HT and FFN206. TLC analysis of VMAT1*_OV_* INS-1 cells preincubated with (A) 20 µM 5-HTP and then supplied for 15 min with Chloroquine 20 µM; Propranolol 20 µM; Fluoxetine 50 µM; Imipramine 100 µM; Reserpine 2 µM; BafilomycinA 200nM. (B) The quantification of 5-HT (n=4) (C) TLC analysis of VMAT1*_OV_* INS-1 cells preincubated with 2 µM FFN206 and then supplied for 15 min with Fluoxetine 50 µM; Reserpine 2 µM; BafilomycinA 200nM. (D) The quantification of FFN206 content (n=4). (E) Schematic representation of the experiment. The cells were treated with FFN206, Chloroquine or 5-HTP and then with BafilomycinA to induce alkalization of the pH in the intracellular compartments. (F) TLC analysis and (G) quantification (n=4; chloroquine n=3) of cellular contents of FFN206, chloroquine and 5-HT after BafA (200 nM) treatment.

**Fig 6.**
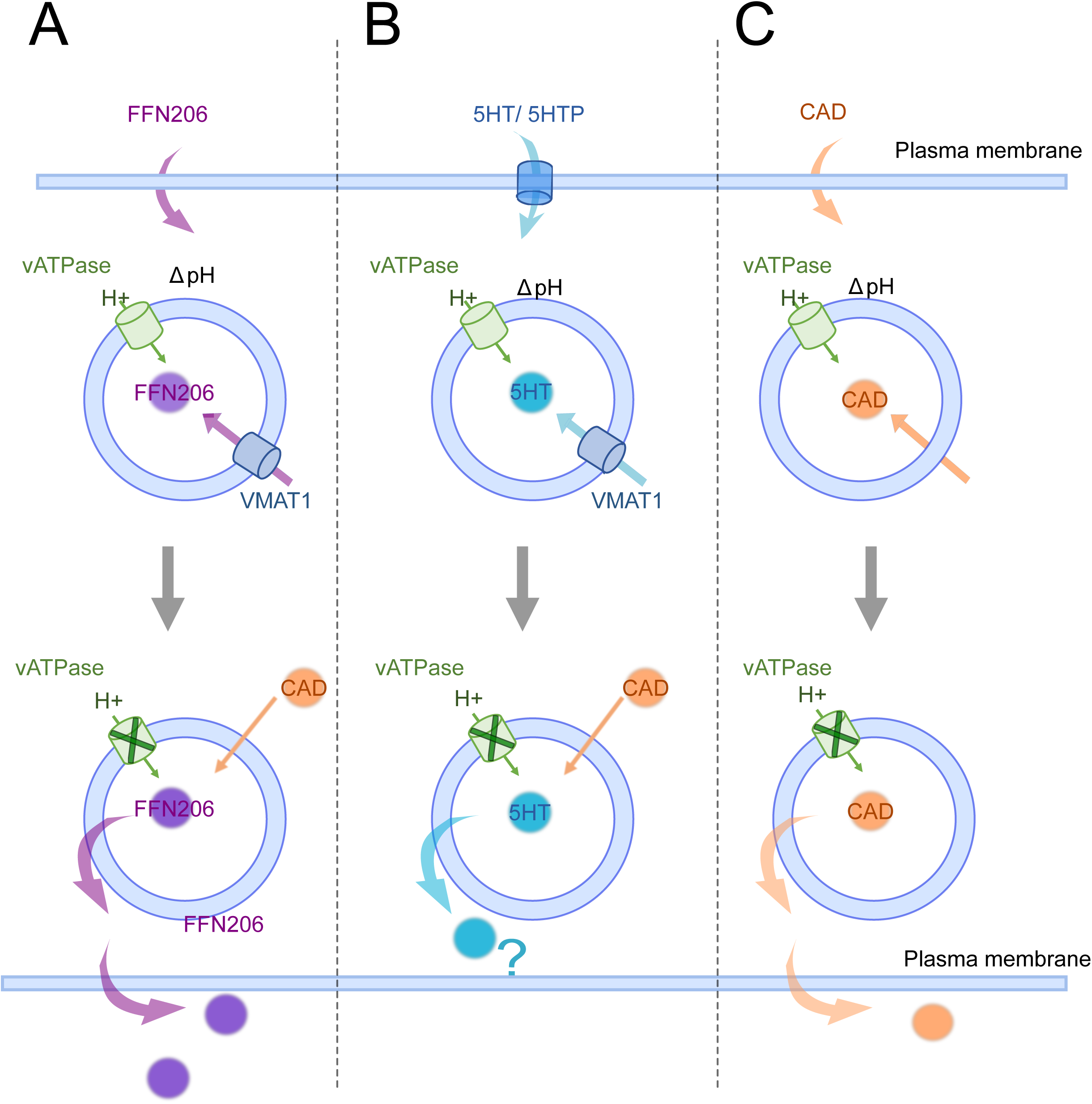
Schematic summary of the results. (A) The artificial monoamine probe FFN206 can move across the plasma membrane (Hu *et al*., 2013). Inside the cell, VMAT1 sequesters FFN206 into SGs. This accumulation is VMAT1-specific and requires a pH gradient, maintained by vATPase. Neuroactive CADs such as propranolol, fluoxetine, imipramine, and clozapine inhibit FFN206 accumulation. and induce the efflux of pre-accumulated FFN206 to the extracellular space. (B) Natural monoamines like 5-HT or 5-HTP require plasma membrane transporters to cross the plasma membrane. 5-HT accumulation in SG of INS-1 cells is VMAT1-dependent (Andersen *et al*, 2016; Hasenhuetl *et al*, 2015; Karkkainen *et al*, 2021).Unlike FFN206, 5-HT is not depleted from cells following pH dissipation or CAD treatment. It is unclear whether 5-HT remains within SGs or is redistributed to the cytosol under these conditions. (C) Lastly, a lipophilic weak base Chloroquine does not require any transporters to cross the membranes and both Its accumulation and retention in cells is pH-dependent.

As observed, CAD, reserpine or pH dissipation with BafA had only limited effect on cellular 5-HT levels, whereas FFN206 signal was strongly reduced (Fig. 5A-D).

We hypothesized that this difference in the efflux is associated with higher membrane permeability of FFN206 than those of 5-HT. To verify this, we compared the cellular retention of FFN206, 5-HT and CQ (membrane permeable CAD), upon pH dissipation with BafA (Fig. 5E). As a result, BafA treatment led to a pronounced loss of both CQ and FFN206, while 5-HT still remained detectable in cell extracts (Fig. 5F, G).

In summary, although FFN206 does not fully recapitulate the behavior of naturally occurring monoamines, it reliably and specifically enables monitoring of VMAT1 activity in INS-1 cells.

## Discussion

This study demonstrates that insulin SGs accumulate both natural monoamine neurotransmitters by VMAT1-facilitated sequestration and synthetic neuroactive drugs through pH-dependent trapping. While these mechanisms operate independently, they converge to influence the SG microenvironment.

Monoamines play established roles in islet physiology, with serotonin promoting insulin secretion and β-cell proliferation during increased metabolic demand (Kim *et al*., 2010; Kim *et al*., 2015; Roberts *et al*., 2023), and dopamine acting predominantly as an inhibitory signal for insulin secretion (Aslanoglou *et al*., 2021; Farino *et al*., 2020). In INS-1 cells, we show that VMAT1 is solely required for vesicular monoamine uptake and for maintaining intracellular serotonin levels (Fig. 1H). Genetic deletion of *Slc18a1* abolished FFN206 accumulation and reduced endogenous 5-HT levels (Fig. 1K and 1H), whereas its re-expression restored vesicular uptake (Fig. 1L). Among monoamines, serotonin was the only neurotransmitter consistently detected, in line with expression of the serotonin biosynthetic machinery in rodent β-cells (Goyvaerts *et al*, 2022; Kim *et al*., 2010). Disruption of monoamine storage by CAD accumulation or VMAT inhibition may therefore influence monoamine-dependent regulatory processes in pancreatic β-cells. Neuropsychiatric disorders and their pharmacological treatments—particularly antipsychotics and antidepressants—are associated with an increased risk of diabetes (Ballon *et al*., 2014; Lee *et al*., 2023; Lindekilde *et al*., 2022). While these associations are multifactorial and largely attributed to central and systemic effects, our findings raise the possibility of an additional peripheral, cell-autonomous contribution. Specifically, neuroactive drugs may accumulate in insulin SGs, altering monoamine handling, and thereby modulating regulated hormone secretion.

Functionally, CADs inhibited VMAT-mediated uptake of the fluorescent monoamine probe FFN206 (Fig.3A, B) and induced rapid efflux of pre-accumulated FFN206 to the extracellular space (Fig.3C-F). This behavior resembles neurotransmitter redistribution described for psychostimulants in synaptic vesicles and chromaffin granules (Freyberg *et al*., 2016; Partilla *et al*., 2006; Rudnick & Wall, 1992b; Sulzer & Rayport, 1990). Notably, CAD-induced redistribution occurred without detectable changes in SG luminal pH (Fig. 4C). By contrast, natural VMAT substrates such as serotonin and dopamine increased the SG pH in a VMAT-dependent manner (Fig 4A, B), consistent with proton–substrate exchange during vesicular loading (Freyberg *et al*., 2016; Sulzer & Rayport, 1990).

Importantly, both inhibition of VMAT-mediated uptake (Fig. 3A, B) and redistribution of pre-accumulated substrates (Fig. 3C-F) are expected to reduce vesicular monoamine content over time. However, the natural VMAT substrate serotonin exhibited substantially greater cellular retention than the fluorescent probe FFN206 under comparable conditions (Fig.5A, B). Previous studies in chromaffin cells and neurons have shown that serotonin efflux from secretory vesicles requires functional plasma membrane serotonin transporters (SERT), whereas VMAT primarily mediates vesicular loading (Freyberg *et al*., 2016; Maron *et al*, 1983; Roz & Rehavi, 2004; Rudnick & Wall, 1992a, b; Sulzer, 2011). Accordingly, limited SERT expression or activity in INS-1 cells may restrict serotonin efflux following vesicular release, in contrast to the more membrane-permeable FFN206. While we cannot directly resolve whether serotonin is released from SGs and retained in the cytosol or remains vesicular upon CAD treatment, its persistence within cells—even after luminal pH dissipation—supports the idea that transporter availability and physicochemical properties jointly shape post-vesicular redistribution. Consistent with this interpretation, chloroquine, like FFN206, was efficiently lost from cells following pH dissipation (Fig.5F, G), in line with its high membrane permeability (Mack & Bonisch, 1979).

In addition to CAD-induced redistribution, we examined the intrinsic retention of FFN206, serotonin, and chloroquine following its withdrawal from the extracellular medium, in the absence of any pharmacological perturbation (EV5A). Under these conditions, serotonin exhibited greater cellular retention than FFN206, consistent with its behavior following CAD treatment, whereas chloroquine showed the highest persistence, remaining detectable for more than 24 hours after withdrawal. These observations further support the idea that post-vesicular retention is strongly influenced by physicochemical properties, particularly membrane permeability.

A relevant experimental consideration is the difference in substrate concentrations used in these assays (2 µM FFN206 versus 20 µM 5-HTP), despite comparable reported Km values for VMAT (∼1–6 µM for both substrates) (Hu *et al*., 2013; Peter *et al*, 1995). Such differences may contribute to distinct retention kinetics. Taken together, these findings indicate that FFN206, while a robust probe for visualizing VMAT-dependent uptake (Hu *et al*., 2013; Yaffe *et al*., 2016), does not fully recapitulate the intracellular dynamics of natural monoamines.

A further mechanistic distinction emerged from the analysis of the SG pH. Natural VMAT substrates, including serotonin and dopamine, increased the SG pH in a VMAT-dependent manner (Fig.4A, B), consistent with previous reports showing alkalinization of synaptic vesicles by amphetamine, MPP⁺, FFN206, and dopamine (Freyberg *et al*., 2016; Sulzer & Rayport, 1990). In contrast, CADs did not measurably alter the SG pH despite inducing FFN206 redistribution (Fig. 4C and Fig. 3C-F). This argues against sustained luminal alkalinization as the primary driver of CAD-induced efflux. Nevertheless, since CQ also did not lead to measurable SG pH rise, we speculate that possible pH alterations could be short-termed and reversible, therefore not detected by the used approach. This would be in line with earlier observations that weak-base-induced alkalinization of endocytic vesicles is transient or heterogeneous (Maxfield, 1982; Sulzer & Rayport, 1990).

In addition, CQ retention in cells is pH-dependent (Fig.5F-G) as dissipation of the proton gradient by BafA efficiently promoted chloroquine loss. Meanwhile, none of the CAD treatments reproduced this effect (Fig. 5F-G; EV5B), supporting the observation that CADs are unlikely to considerably change the SG pH.

Together, our results highlight a fundamental difference between *bona fide* VMAT substrates and CADs: substrates rely on vesicular transport and can directly influence both SG content and pH, whereas CADs accumulate independently of active transport, alter vesicular content without sustained pH changes, and are governed primarily by membrane permeability.

Several limitations should be acknowledged. This study relies on INS-1 cells expressing VMAT1, whereas human β-cells predominantly express VMAT2 (Schafer *et al*., 2013). While both isoforms share conserved transport mechanisms (Finn *et al*, 1998), future studies in human β-cells will be required to assess quantitative differences. In addition, CAD accumulation in insulin SGs is inferred from functional assays rather than direct ultrastructural visualization, and in vivo accumulation of these drugs in pancreatic islets remains to be shown. Future studies in primary human tissue and animal models will be required to determine the physiological relevance of these mechanisms under pharmacological exposure.

In summary, insulin SGs emerge as acidic organelles capable of accumulating cationic amphiphilic drugs. By distinguishing VMAT-dependent monoamine loading from pH-driven trapping of synthetic weak bases, this work uncovers distinct mechanisms through which endogenous metabolites and pharmacological agents affect SG homeostasis. These principles are likely to extend to large dense-core vesicles in other peptide hormone-secreting endocrine cells and neurons.

## Data availability

The Python code used for the segmentation, FLIM analysis, and SG pH interpolation is available on GitHub (https://github.com/olekstop/flim_granules_analysis)

## Author contributions

OT, MS: study concept. OT: design and performance of all experiments with technical assistance by AS, KG, CW, and CM. OT, MLZ: writing of the Python code for FLIM. OT, AM: SIM. MN, JT, K-PK: generation of constructs. MN: generation of *Slc18a1^−/−^* clones. OT, ST: LC-MS/MS for CADs. OT, MG: liposome studies. OT, MS, MG, AM: experimental design and data interpretation. All authors discussed the results. OT: wrote the first draft of the manuscript. OT, MG, MS: editing of the manuscript with input from all other authors.

## Acknowledgements

We wish to thank Maria Federova, Palina Nepachalovich, Iuliia Iermak, Ünal Coskun Johannes Broichhagen, Stefan Diez, and Graem Eisenhofer for discussion; Mirko Peitzsch for initial measurement of monoamines by LC-MS/MS. Sider Penkov and Cornelia Wetzker for advice about TLC and FLIM, respectively; Moeko Sakate and Obada Bahra for experimental help. We are grateful to the colleagues of the Light Microscopy Facility of the CRTD at TU Dresden and the technicians of the Institute of Clinical Chemistry of the University Hospital Dresden for their support. We are also grateful to Katja Pfriem and Aline Kalinka for administrative help. These studies were supported with funds from the German Center for Diabetes Research (DZD e.V.) by the German Ministry for Research, Technology and Space (BMBFTR); from the Innovative Medicines Initiative 2 Joint Undertaking under grant agreement No 115797 (INNODIA) and No 945268 (INNODIA HARVEST). This Joint Undertaking received support from the European Union’s Horizon 2020 research and innovation program, European Federation of Pharmaceutical Industries and Associations, JDRF (now Breakthrough T1D), and The Leona M and Harry B Helmsley Charitable Trust; the European Union’s Horizon Europe research and innovation program under grant agreement No 1010954433. OT has been the recipient of a DZD Outstanding Young Investigator Award 2025 and was supported by a travel allowance from the DIGS-ILS mentoring program. PT has been the recipient of a visiting fellowship from TU Dresden and an EFSD Albert Renold Travel Fellowship.

*Funded by the European Union. Views and opinions expressed are, however, those of the author(s) only and do not necessarily reflect those of the European Union. Neither the European Union nor the granting authority can be held responsible for them.*

## Materials and Methods

### Cell culture

INS-1 cells were cultured in RPMI 1640 medium supplemented with 10% fetal bovine serum (FBS), 100 U/mL penicillin, 100 μg/mL streptomycin, 2 mM L-glutamine, 50 μM β-mercaptoethanol, 1 mM sodium pyruvate, 10 mM HEPES pH 7.4. Cells were maintained at 37°C in a humidified atmosphere with 5% CO₂. Transient transfection was performed with Amaxa Nucleofector™ II (Lonza) using the T-20 program and transfection reagents according to the manufacturer’s instructions.

### Immunocytochemistry

INS-1 cells were grown on high-precision coverslips coated with poly-L-ornithine. Cells were fixed with 4% paraformaldehyde (PFA) and permeabilized with 0.3 % Triton X-100. After 1 hr blocking with 0.2% fish skin gelatin, 0.5% BSA in PBS, cells were incubated with primary antibodies for 2 hrs at RT (1:200) in a humidified chamber and washed 3x with PBS. Secondary antibodies were kept for 30 min at RT (1:200). Following 3x PBS wash, nuclei were stained for 10 min with DAPI (1:5000). After a final wash, coverslips were mounted on a glass slide with Mowiol (Calbiochem).

### Confocal imaging and structured illumination microscopy (SIM)

Confocal microscopy or structured illumination microscopy (SIM) images were acquired with Nikon C2+ confocal and SIM-E microscope with a Nikon Plan Apo λ 60x Oil objective or for SIM 100x Oil objective with a Hamamatsu camera (for SIM). The confocal system is equipped with lasers with emission wavelengths of 405, 488, 561 and 640 nm. The wavelengths of the SIM lasers are 488, 561 and 640 nm. The collected SIM images were reconstructed using the Nikon NIS Elements software, and then contrast and brightness levels were adjusted using the FIJI software (National Institutes of Health) (Schindelin *et al*, 2012).

### Imaging of FFN206 by confocal fluorescence microscopy

To study the accumulation of FFN206 (Tocris), cells were plated onto poly-L-ornithine coated, 8-well µ-slides (Ibidi). To study the influence of compounds, the culture medium was exchanged to media containing the compound in water/ DMSO or vehicle alone (only for Clozapine), and incubated for 40 min. After that, 2 μM FFN206 was added for 1 hr Prior imaging media was replaced with imaging buffer (HEPES 15 mM, KCl 5 mM, Glucose 11 mM, NaCl 120 mM, NaHCO_3_ 24 mM, MgCl_2_ 1 mM, CaCl_2_ 2 mM, Albumin 1 mg/mL, pH 7.4). Fluorescence images (3-5 images/condition) were acquired with a Nikon C2+ confocal microscope equipped with a Nikon Plan Apo λ 60x Oil objective.

The FFN206 was excited with the 405 laser; emission was set to standard DAPI settings. All images were adjusted using the same contrast and brightness level using FIJI ImageJ software (National Institutes of Health) (Schindelin *et al*., 2012).

For testing the efflux of FFN206, cells were stained with 2 μM FFN206 for 1 hr in culture medium, which was then replaced with the imaging buffer. Compounds at concentrations indicated in the figure legend were spiked into it. Cells were imaged directly after the addition of the compound with the same settings as the non-treated control.

### FFN206 accumulation in a 96-well plate format

The FFN206 uptake assay was adapted from (Hu *et al*., 2013) *Slc18a1*^−/−^ INS-1 cells were transfected with either VMAT1 or an empty vector (pcDNA3.1) and grown in 96-well poly-L-ornithine-coated black plates. The fluorescence was measured using a BioTek Synergy H1 multi-mode plate reader (excitation 369 nm, emission 464 nm). For the analysis, nonspecific uptake, measured in pcDNA3.1-transfected cells, was subtracted from the FFN206 signal measured in VMAT1-transfected cells. The data were normalized to the untreated control, and results are presented as a percentage of the untreated sample. Two *Slc18a1*^−/−^ INS-1 clonal cell lines were analyzed separately with individual controls for each.

### Measurements of SG luminal pH

Fluorescence lifetime imaging microscopy (FLIM) based measurements of pH were performed as previously described by (Neukam *et al*., 2017) (preprint). When applicable, cells were treated with compounds, added directly to the imaging buffer, and incubated for 15 min prior to imaging. In the case of longer incubation time (for treatment with 5-HTP and L-DOPA for 1 hr), cells were maintained in full growth media spiked with compounds, which were replaced with the imaging buffer (details about time and concentrations are given in the figure legends). During imaging, cells were maintained at 37°C in the imaging buffer. FLIM was performed using a Leica SP8 upright confocal microscope with an HC APO UVIS CS2 63x/0.9 water objective. The lifetime of eCFP was recorded using a multiphoton laser set to 880 nm with 80 MHz laser frequency and 12.5-time interval between pulses, and photons were detected in counting mode with emission set to 462-530 nm. FLIM images were acquired over 25 frames with an acquisition of 400 µs per frame. Laser intensity was optimized to avoid pile-up, achieving <1 photon per pulse. In each experiment, 4-5 images per condition were acquired, and for each condition, at least three independent measurements on separate days were performed. The images were fitted with a second-order exponential decay model, using the n-Exponential Reconvolution algorithm in the Leica Application Suite X (LAS X, version 3.5.7.23225). Instrument response function (IRF) background and shift were corrected with binning set to 1 and a photon threshold of 50 counts. For further analysis, fitted images were saved in TIFF format with scaling factors for the lifetime of 0.001 per gray level (to convert nano to picoseconds) and 0.01 for chi-square values. The exported data included photon counts (intensity images), amplitude-weighted lifetime, and chi-square maps to evaluate fit quality. The detailed analysis pipeline, including the model as well as the full Python code, is described: (*flim_granules_analysis*; GitHub)

### Extraction and thin-layer chromatography (TLC)

For all experiments, cells were grown on poly-l-ornithine-coated wells (0.1 mg/mL). For compound accumulation, fresh medium containing the compounds was added for 1 hr. Then, medium was aspirated, cells washed 3x with warm DPBS, and harvested by scraping in 1 mL DPBS following cell pelleting. The pellets were either directly extracted or snap frozen and stored at −80°C.

For extraction from cells, cell pellets were resuspended in 150 µL of extraction solvent (CH_3_CN:MeOH:H_2_O (40:40:20, v/v)+ 0.1% HCOOH) and lysed 30 minutes on ice. Protein precipitates were removed by centrifugation at 12,000 x *g*, for 30 min at 4°C. Supernatants were vacuum-dried and dissolved in 15 µL of the same solvent. 3 µL of the sample was applied onto the HPTLC plates and developed in a single pass in CHCl₂:MeOH:Me2CO:AcOH₂O (46:15:17:14:8 v/v). After, the plates were dried and visualized with UV light.

For compound extraction from the growth media, 500 µL was alkalized by adding 32 µL of 3 M NaOH (to reach final pH∼13) and subjected to biphasic extraction with ethyl acetate (2:1; 1 mL of ethyl acetate was added to 500 µL of media, followed by vortexing and shaking at 1,200 rpm for 7 minutes). The samples were centrifuged at 5,000 x *g* for 30 min, and the organic phase was collected into a separate tube. The bottom phase was re-extracted, and the combined organic extracts were dried.

The fluorescent signal from TLC was normalized using cholesterol content in the extracted sample, which was measured by Amplex Red Cholesterol Assay Kit (Invitrogen). Cell extracts (3 ul of the same that were applied on the TLC) were mixed with 1x Reaction buffer from the Amplex Red Cholesterol Assay Kit, and the assay was further performed according to the manufacturer’s instructions.

### Retention (efflux) assay in INS-1 cells

Cells were incubated for 1 hr in the presence of the respective compound for accumulation. Afterwards, a counter compound was spiked directly into the 1 mL of media without any washing step and incubated for 15 or 60 min, as indicated in the figure legend. Post-treatment, cells were washed with 2x DPBS and extracted as described above.

For the efflux experiments, where growth media were analyzed, after 1 hr of accumulation of 2 µM FFN206, cells were washed 3x with warm DPBS. After, 600 µL of fresh media containing the chosen compound was added and kept for an additional 15 min to allow for the redistribution of pre-accumulated FFN206. After this incubation, 500 µL of media was collected in separate tubes. Then, cells were washed with warm DPBS and harvested for extraction as described before.

### Accumulation of CADs in liposomes

Liposomes were prepared using POPC (1-palmitoyl-2-oleoyl-sn-glycero-3-phosphocholine) and cholesterol at a molar ratio of 70:30. The organic solvents were evaporated under a gentle stream of nitrogen. The dried lipid film was rehydrated in either neutral (25 mM HEPES, 150 mM KCl, pH 7.5) or acidic buffer (25 mM citrate, 150 mM KCl, pH 5.5) to reach a final lipid concentration of 2.5 mg/mL. The resulting multilamellar vesicles were subjected to extrusion through a 100 nm polycarbonate membrane filter at 60°C. The suspension was passed 15 times through the membrane until a homogeneous, non-turbid solution was obtained. Liposome size and polydispersity were assessed by dynamic light scattering (DLS) with Zetasizer Ultra (Malvern). The liposomes were then aliquoted, snap-frozen, and stored at −80°C until further use.

On the day of the experiment, liposomes were diluted to a concentration of 1.25 mg/mL. Next, the liposomal buffer was exchanged using PD MiniTrap G-10 desalting columns (Cytiva) equilibrated with either neutral or acidic buffer, which gave three types of liposomes: (a) Liposomes in Neutral buffer and containing acidic buffer in the lumen (pH gradient); (b) Liposomes in an acidic buffer and containing an acidic buffer in the lumen (no pH gradient); (c) Liposomes in Neutral buffer and containing Neutral buffer in the lumen (no pH gradient). After buffer exchange, 150 µg of liposomes were incubated with respective compounds at a final concentration of 40 µM for 2 hrs at RT. Control samples, including liposomes alone and compounds alone, were processed in parallel. Following incubation, the reaction was diluted with 1.1 mL of the corresponding outer buffer. The samples were then transferred to Beckman ultracentrifuge tubes and ultracentrifuged with TLA 55 Fixed-Angle Rotor (Beckman Coulter) at 40,000 rpm for 1 hr at 4°C. The supernatant was carefully aspirated, and the liposome pellet was resuspended in 100 µL of Milli-Q water and transferred to a fresh tube. The liposome sample was alkalinized with 3 µL of 25% ammonia solution (pH∼12-13). The samples were then spiked with 15 µg of DOPC (1,2-dioleoyl-sn-glycero-3-phosphocholine) and 4 µM Clozapine, which were used as a lipid and drug internal standards, respectively. The sample, including internal standards, was then extracted with three volumes of MeOH:CHCl_3_ (1:10, v/v). The sample was vigorously vortexed, spinned 5,000 x *g* for 5 min, at 4°C and the organic (lower) phase was collected. The upper phase was re-extracted in a similar way, and the two consecutive extracts were pooled and dried under vacuum. The dried extract was resuspended in 100 µL of methanol containing 0.01% HCOOH. Samples were analyzed using direct injection with an Advion compact mass spectrometer (expression®L CMS) operated in positive ion mode with ESI at standard instrumental settings. The m/z detection ranges were set to 250–350 for drug compounds and 750–850 for lipids. For data analysis, the peak area of each compound or lipid was obtained and quantified using the respective internal standard. Drug measurements were quantified using clozapine as an internal standard and further normalized to the POPC content in the extracts (quantified by DOPC as internal standard) (EV3).

### LC-MS/MS of endogenous monoamines

Cells from a T75cm^2^ flask were collected by harvesting in 10 mM EDTA in PBS, centrifuged 1,200 x *g* for 5 min at 4°C and re-suspended in 120 µL of Milli-Q water. 20 µL of the resulting cell mixture was taken for total protein quantification with the Pierce™ BCA Protein Assay Kit (Thermo Fisher Scientific). The rest was extracted with 500 µL of CH_3_CN:MeOH:H_2_O (40:40:20, v/v) + 0.1% HCOOH. The proteins were precipitated for 30 min on ice, following centrifugation 16,000 x *g* for 30 min.

Separation was performed on an ultra-performance liquid chromatography system equipped with a Phenomenex Kinetex (2.6 µm Biphenyl 100×2.1). Analytes were separated by gradient elution at 40°C with a flow rate of 0.3 mL/min. The gradient was set as follows: (i) 0–0.1 min: 10% B, (ii) 0.1–1.5 min: 10-95 % B, (iii) 1.5-2 min: 95% B (isocratic, column wash), (iv) 2–2.3 min: 95-10% B, (v) 2.3–3.8 min: 90% B (isocratic, column re-equilibration). Mobile phase composition, A: H_2_O + 0.1% HCOOH; B: CH_3_OH and 0.1% HCOOH.

Mass spectrometry was conducted in multiple reaction monitoring scan mode (MRM) using positive electrospray ionization including the ion source parameters: curtain gas (35 psi), collision gas 9 (psi), ESI voltage (4,500V), source temperature (600°C), gas 1 (50 psi) and gas 2 (50 psi). Optimized parameters are provided in EV Table 3.

### LC-MS/MS analysis of CADs

One experimental condition (one sample) was extracted from ∼up to 100 µg of total protein by BCA (as described earlier). Before extraction, cell pellets were spiked with 10 µL of two separate internal standard mixes. First, a mix containing Clozapine-D4, Propranolol-D7, and Fluoxetine-D6 was prepared in MeOH with a final amount of each standard 10 pmol per sample. For 5-HT and DA quantification, a separate internal standard mix was prepared in 100 mM CH_3_COOH. The final concentration of spike was 56.7 pmol 5-HT and 65.2 pmol dopamine-D4. Next, the proteins were precipitated with 150 µL of *i*-PrOH + 0.1% HCOOH and incubated on ice for 30 min with occasional vortexing and centrifuged at 13,000 x *g* for 10 min. Cleared supernatant was collected for biphasic extraction and 350 µL of MTBE:MeOH 10:3 (v/v) + 0.1% HCOOH was added to the supernatant. The mixture was agitated at 1200 rpm for 1 hr at 4°C. Phase separation was induced by adding 300 µL of water, followed by shaking at 1200 rpm for 15 min. The samples were then centrifuged at 13,000 x *g*, for 15 min, 4°C, and the lower aqueous phase was carefully collected and dried under vacuum. Samples were reconstituted in 100µL of 1:1 (A:B) mobile phase before injection. Mobile phase composition as follows, A: H_2_O + 10 mM HCOONH_4_and 0.1% HCOOH; B: CH_3_CN:H_2_O (90:8.9, v/v) + 10 mM HCOONH_4_ and 0.1% HCOOH.

Analysis of compounds was done by liquid chromatography tandem mass spectrometry (LC-MS/MS). Separation of the analytes was performed on an ultra-performance liquid chromatography system (Shimadzu) equipped with a C18 CORTECS Waters column (2.7 µm, 2.1 × 100 mm, Waters). 10 µL of the sample (or calibrator) was injected onto the column and separated by gradient elution at 40°C with a flow rate of 0.3 mL/min. The gradient was set as follows: (i) 0–13 min: 5-75% B, (ii) 13–13.1 min: 75-100% B, (iii) 13.01-17 min: 100% B (isocratic, column wash), (iv) 17–17.1 min: 100-5% B, (v) 17–22 min: 5% B (isocratic, column re-equilibration).

Mass spectrometry was performed with triple quadrupole linear ion trap mass spectrometer Triple Quad 7500 (SCIEX, Framingham, MA, USA) in multiple reaction monitoring scan mode (MRM) using positive electrospray ionization including the ion source parameters: curtain gas (40 psi), ESI voltage (5,500V), source temperature (500°C), gas 1 (70 psi) and gas 2 (50 psi). The optimization of MRM parameters was carried out by direct injection of each analyte / isotopically labelled standard (10 µL/min), diluted in CH_3_CN:H_2_O (50:50, v/v). 2-3 fragments per analyte was selected based on the sensitivity. The parental ion and selected product ions, along with optimized fragmentation parameters are provided in EV Table 4.

MS data was analyzed using Skyline (v24.1 by MacCoss Lab Software) and quantified by isotopically labeled internal standards. CQ and imipramine were quantified by Clozapine-D4 (due to the unavailability of ring-labelled corresponding standards). The data were normalized to total protein, measured by BCA, as described earlier.

### Statistics and software

The normality was verified with the Shapiro-Wilk test. For normally distributed data, statistical significance was determined using one-way analysis of variance (ANOVA; α = 0.05) in OriginLab 2024b (Academic). For statistical analysis of FLIM data and normality of distributions was assessed using the Shapiro-Wilk test, and non-parametric comparisons were performed using the Mann-Whitney U test. Data manipulation and statistical testing were performed using the pandas and scipy libraries in Python.The p values on the plots are denoted as: > 0.05 n.s (asterisk not shown); ≤ 0,05 *; ≤ 0,01 **; ≤ 0,001 ***; ≤ 0,0001 ****. Graphs were generated using OriginLab 2024b (Academic) or Python (mainly with the seaborn library) for FLIM data visualization. Image analysis was performed using Fiji (ImageJ) (Schindelin *et al*., 2012) and Python (napari) (Sofroniew, N. (2025). The MS data, after acquisition with SCIEX operating software, were analyzed in Skyline (v24.1 by MacCoss Lab Software) (MacLean *et al*, 2010). Chemical structures were generated with ChemSketch (ACD/Labs). For the physicochemical properties of the drugs and metabolites, the web resources PubChem and HMDB were used unless a specific reference is indicated in the text.

